# PD-L1 deletion or blockade regulate macrophage antigen presentation and checkpoint molecule surface levels

**DOI:** 10.64898/2026.06.23.734016

**Authors:** Trinity Waddell, Haidong Dong, Minna Roh-Johnson, Jessica N. Lancaster

## Abstract

Macrophages in the tumor microenvironment are known to upregulate PD-L1 expression, thereby suppressing T cells through PD-1 ligation. However, the manner in which PD-L1 expression intrinsically impacts macrophages and their immunomodulatory phenotype is less clear. Clarifying this knowledge gap would yield insight into the mechanisms of immunosuppression within the tumor microenvironment. To characterize the macrophage intrinsic role of PD-L1, we used complementary genetic and pharmacological approaches by analyzing primary murine bone marrow-derived macrophages (BMDMs) with complete genetic PD-L1 deletion and wildtype BMDMs treated with anti-PD-L1 blocking antibodies. Macrophages were evaluated across naïve, pro-inflammatory (M1), and tumor conditioned (TCM) polarization states in vitro. Unlike prior reports, neither genetic deletion nor antibody blockade dramatically altered the expression of macrophage polarization markers or in vitro phagocytic capacity. Both conditions consistently reduced surface levels of the M1-associated costimulatory molecule CD80, prompting further analysis of T cell interacting and antigen presenting proteins, in which we revealed disparate effects of genetic deletion and antibody blockade on the surface levels of MHCI, MHCII, PD-1, and PD-L2. These findings suggest that PD-L1 deletion and antibody-mediated blockade contribute to macrophage immune regulatory profiles in distinct manners. This difference supports a model in which PD-L1 functions in macrophages beyond its canonical role as a ligand for PD-1, influencing antigen presentation and checkpoint molecule levels and playing a broader role in immune regulation in the tumor microenvironment.

## Introduction

Strategies to successfully target tumor macrophages are needed to improve patient therapeutic responses. Macrophages are highly plastic innate immune cells that play an essential role in shaping the tumor microenvironment. They display a spectrum of polarization states, classically defined by a framework of pro- versus anti-inflammatory states, by which they facilitate myriad roles in infection, cancer, and tissue homeostasis (1–3). Cell-extrinsic and - intrinsic cues shape macrophage polarization states, which, in turn, enforce the context-specific activation states of T cells. Because of their immunomodulatory functions and their ability to take on unique polarization or functional profiles in response to local microenvironmental cues, macrophages play an important role in the efficacy of immunotherapy (4,5).

The canonical function of programmed cell death-ligand 1 (PD-L1), a membrane-bound immune checkpoint ligand protein, is to bind the T cell inhibitory receptor programmed cell death protein 1 (PD-1), thereby suppressing T cell activation and adaptive immune responses (6–8). PD-L1 is frequently overexpressed on cancer cells as a mechanism of immune evasion, leading to suppression of T cell activation in the tumor microenvironment (9,10). Immune checkpoint blockade therapies have been developed to interfere with the PD-1/PD-L1 axis with blocking antibodies to restore cytotoxic T cell function (11,12). This therapeutic has had significant impact on patient outcomes in many types of cancers, underscoring the clinical relevance of PD-L1 (13).

While PD-L1 overexpression in cancer cells has been a primary target in PD-1/PD-L1 immune checkpoint blockade, PD-L1 expression by tumor-associated macrophages also plays a significant role in therapeutic outcomes (14–16). Prior work has indicated potential differential efficacies of PD-1 and PD-L1 blocking antibodies (17–20), suggesting that the cellular mechanisms driving outcomes of each monotherapy may differ. Despite its limited cytoplasmic domain, PD-L1 has been shown to have cell-intrinsic functions independent of PD-1 ligation, including intracellular and nuclear activities in a variety of cancer cell types (21,22) and in dendritic cells (23). Connections between PD-L1 expression and macrophage activation status within tumors have also been noted (24). Subsequent studies in our lab (25) and others (17,26–28) suggested that treatment of macrophages in monoculture with PD-L1 blocking antibodies promoted an immunostimulatory phenotype and function. These findings suggest that PD-L1 exerts function independent of the canonical PD-1/PD-L1 axis. However, the literature remains inconsistent as to whether PD-L1 expression intrinsically promotes or stabilizes macrophage states.

There currently remains a lack of understanding as to whether PD-L1 actively regulates macrophage activation or merely serves as a marker of macrophage activation state. PD-L1 expression is highest in M1 inflammatory macrophages and lowest in naïve macrophages (29). This is likely due to its upregulation induced by interferon-gamma (IFNγ), (30–32) but PD-L1 is also elevated in tumor conditioned macrophages over the naïve state. Because of the evidence that macrophage PD-L1 expression strongly influences the tumor immune environment, it is interesting to note the disparity in expression between inflammatory M1 and tumor conditioned macrophages. Though there has been some evidence of association, it is not yet clear whether PD-L1 influences macrophage polarization and ability to impact the immune microenvironment. This points to a critical need to better characterize PD-L1’s intrinsic roles in modulating macrophage functions independent of PD-1 interaction. Our goal was to address this using genetic and pharmacological approaches. Towards this end, we compared murine macrophages with complete genetic abrogation of PD-L1 expression and those treated by antibody-mediated PD-L1 blockade against PD-L1-sufficient controls. For murine macrophages in isolation, we found that PD-L1 inhibition did not broadly impact macrophage polarization profiles or phagocytic capacities. Our results indicated that abrogation and blockade of PD-L1 induced changes to macrophage expression of antigen presentation and immune checkpoint molecules in distinct ways, supporting a strategy of PD-L1 targeting to modify macrophage interactions within the tumor microenvironment.

## Results

### PD-L1 blockade and genetic abrogation have limited impact on macrophage polarization

We established an in vitro monoculture system in which we differentiated primary murine bone marrow derived macrophages (BMDMs) from C57BL/6J (PD-L1 WT) or B6.B7-H1^-/-^ (PD-L1 KO) mice(30). Naïve BMDMs (M0) were polarized towards a pro-inflammatory (M1) phenotype by the addition of IFNγ and lipopolysaccharide (LPS), or a tumor conditioned (TCM) state using interleukin-4 (IL-4), interleukin-10 (IL-10), and E0771 murine triple negative breast cancer cell line supernatant (**Figure 1A**). For the PD-L1 blockade group WT macrophages were treated with either anti-PD-L1 antibodies (clone 10F.9G2) or isotype controls during the 48-hour polarization step of our experimental workflow (**Figure 1A**). Macrophages were polarized for 48 hours consistent with prior literature (31,32). PD-L1 deficiency and blockade were confirmed by detection of anti-PD-L1 surface staining with flow cytometry, and deficiency was further confirmed by Western blot detection of whole PD-L1 protein from lysates of M1-polarized BMDMs (**Supplementary Figure 1A**).

**Figure 1.**
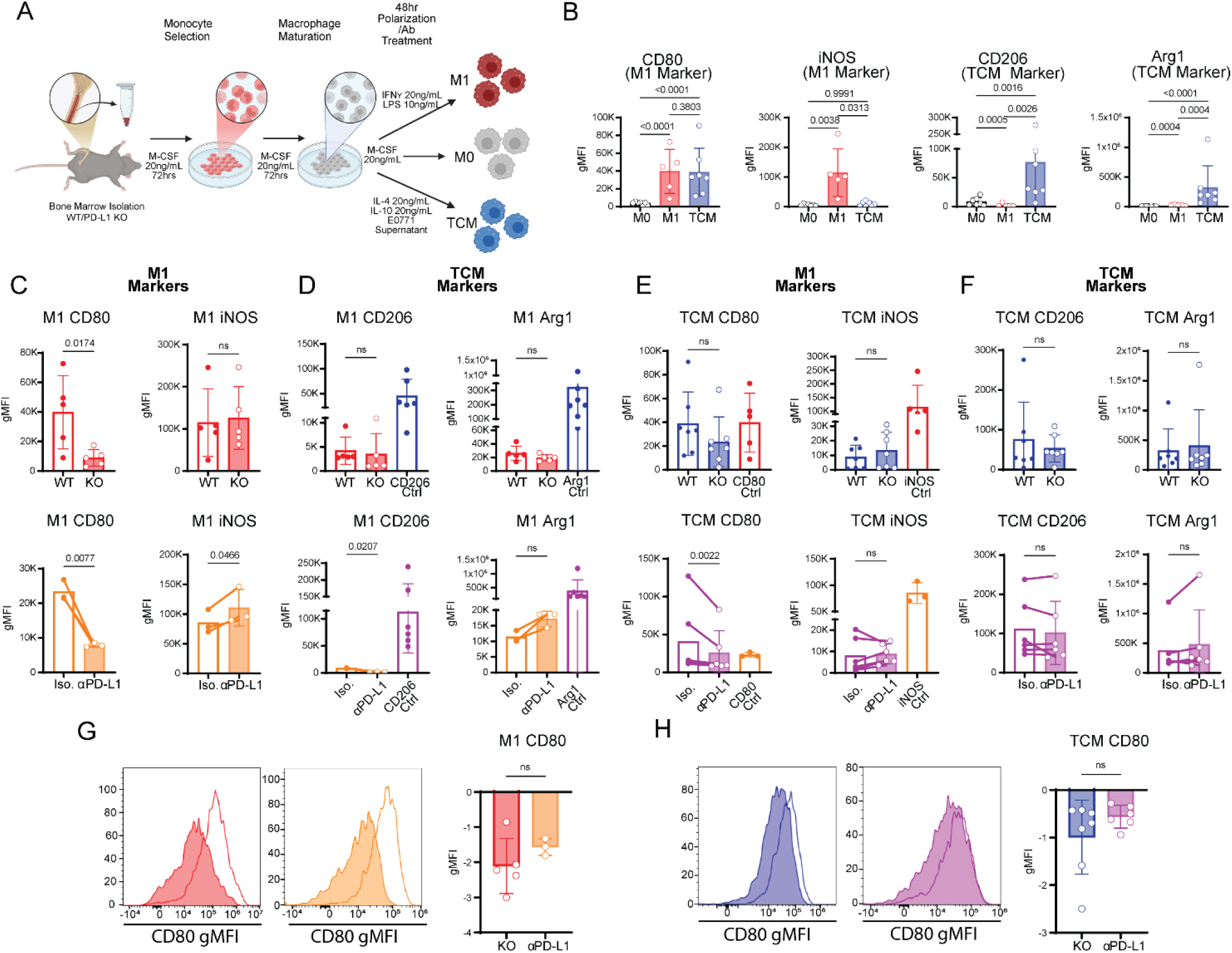
PD-L1 blockade and abrogation have limited impact on macrophage polarization. (A) Experimental setup: Bone marrow cells isolated from PD-L1-sufficient (WT) and -deficient (KO) mice were differentiated into naïve M0 BMDMs using M-CSF and polarized to M1 or TCM states by IFNγ/LPS or IL-4/IL-10/EO771 tumor medium for 48 h. WT Mice were also treated with either anti-PD-L1 antibody or isotype antibody during polarization. Legend for bar graph organization. Colors organized by Polarization: M1 Red and Orange, and TCM Blue and Purple, and by PD-L1 interference: WT/KO Red and Blue, Iso./Blockade Orange and Purple. (B) Flow cytometry Geometric mean fluorescence intensities (gMFI) of M1-associated (CD80, iNOS) and TCM-associated (CD206, Arg1) polarization markers in WT M0, M1, and TCM polarized BMDMs. N = 5-7 mice in separate experiments. Data analysis performed with multiple T-tests and mixed effects with Tukey’s multiple comparisons post-hoc analysis on log10 transformed data. (C) gMFI of M1 markers CD80 and iNOS upon M1 polarization of BMDMs comparing WT vs KO (N=5 mice analyzed in 3 experiments)(CD80 geometric mean fold change = 0.23 (95% CI 0.08 – 0.72)) and Isotype vs Blockade conditions. (N=3 mice analyzed in 1 experiment)(CD80 and iNOS geometric mean fold change = 0.34 (95% CI 0.23 – 0.51), = 1.29 (95% CI 1.01 – 1.64)). (D) gMFI of TCM markers CD206 and Arg1 upon M1 polarization of BMDMs comparing WT vs KO (N=5 mice analyzed in 3 experiments) and Isotype vs Blockade (N=3 mice analyzed in 1 experiment) conditions. (E) gMFI of M1 markers CD80 and iNOS upon TCM polarization of BMDMs comparing WT vs KO (N=7 mice analyzed in 5 experiments) and Isotype vs Blockade (N=6 mice analyzed in 4 experiments) conditions (CD80 geometric mean fold change = 0.68 (95% CI 0.57 – 0.81)). (F) gMFI of TCM markers CD206 and Arg1 upon TCM polarization of BMDMs comparing WT vs KO (N=7 mice analyzed in 5 experiments) and Isotype vs Blockade conditions (N=6 mice analyzed in 4 experiments.). (G) Fold change in CD80 expression of M1 polarized BMDMs comparing KO over WT and Blockade over Isotype treatment. (N=3-5 mice from 1-3 experiments). (H) Fold change in CD80 expression of TCM polarized BMDMs comparing KO over WT and Blockade over Isotype treatment. (N=5-7 mice from 3 experiments). Welch’s corrected T-test used on log10 transformed gMFI to compare WT vs KO. Control columns correspond to the associated polarization state and are shown for biological context, not included in analysis. Paired T-test used on log10 transformed gMFI to compare isotype vs anti-PD-L1 treated. Control columns correspond to the associated polarization state are shown for biological context, not included in analysis. Due to unmixing of fluorescent signals a constant value was used to ensure all samples expressed a positive fluorescent signal. The constant was applied to all samples in the condition and always resulted in the lowest expressing sample having a signal of 1000. A constant of 3282 and 1487.333 were applied to panel D WT/KO CD206 and Iso./aPD-L1 CD206 respectively.

The baseline phenotypes of WT M0 BMDMs and those polarized to M1 and TCM states were characterized using flow cytometry. WT M0 BMDMs expressed low levels of costimulatory factor CD80, inducible nitric oxide synthase (iNOS), macrophage mannose receptor (CD206), and arginase-1 (Arg1) (**Figure 1B**). WT M1 BMDMs highly upregulated CD80 and iNOS, whereas WT TCM BMDMs significantly upregulated CD206 and ARG1 (**Figure 1B**). WT TCM BMDMs also upregulated CD80 (**Figure 1B**). Given these marker level shifts with polarization, CD80 and iNOS were used to define M1 polarization, while CD206 and Arg1 were used to define TCM polarization (3,33,34). The baseline polarization profile of WT BMDMs treated with isotype antibodies was also confirmed by flow cytometry and largely matched the profile of WT BMDMs (35) (**Supplementary Figure 1B**).

The expression of polarization markers was compared between WT and KO BMDMs and isotype and PD-L1 blockade treated BMDMs in M0, M1, and TCM states. M0 macrophages consistently had low surface levels of the markers measured, resulting in no significant differences in the M1 markers CD80 and iNOS (**Supplementary Figure 1C**), or TCM markers CD206 and Arg1 (**Supplementary Figure 1D**). Both PD-L1 KO and PD-L1 blockade M1-polarized BMDMs had significantly decreased CD80 surface levels as compared to WT and isotype treated M1-polarized BMDMs (**Figure 1C**). While there was no change in the M1 marker iNOS in PD-L1 KO BMDMs, there was an increase in iNOS expression upon PD-L1 blockade (**Figure 1C**). TCM markers CD206 and Arg1 were largely unchanged in M1-polarized KO BMDMs compared to WT, though there was a statistically significant decrease in CD206 surface levels upon PD-L1 blockade in M1-polarized macrophages (**Figure 1D**). To provide biological context for the decreased CD206 expression within M1-polarized macrophages, CD206 and Arg1 expression levels were graphed alongside WT TCM macrophage controls. Likewise, WT M1 levels of M1 markers CD80 and iNOS were graphed as controls for TCMs. PD-L1 KO TCM polarized BMDMs showed a trend of a decrease in CD80 surface levels but no change in iNOS expression, while PD-L1 blockade resulted in a significant decrease in CD80 surface levels and no change in iNOS expression (**Figure 1E**). Analysis of TCM markers CD206 and Arg1 in TCM polarized macrophages demonstrated no significant changes upon PD-L1 KO or blockade (**Figure 1F**). Thus far, the most notable impact of PD-L1 KO or blockade was the consistent decrease in CD80 surface levels across conditions.

To confirm that the observed changes in marker levels were not due to differential cell survival across conditions, we analyzed viability of WT, KO, isotype and PD-L1 blockade BMDMs following 48 hours of M1 and TCM polarization. We observed maintained or increased viability with PD-L1 KO or blockade in all conditions. Notably, TCM-polarized KO BMDMs and M1-polarized PD-L1 blockade BMDMs demonstrated significantly improved survival of the stress of polarization relative to their controls (34–36) (**Supplementary Figure 2A**). These differences suggest that the limited changes observed were not due to preferential survival of less polarized macrophages.

To compare the effects of PD-L1 KO versus PD-L1 blockade on macrophage polarization, we compared log_2_ fold changes relative to respective controls for markers with statistically significant differences. For M1-polarized macrophages we observed no difference in fold change of levels of CD80 or iNOS between PD-L1 KO and PD-L1 blockade macrophages (**Figure 1G**). While there was a significant difference between KO and anti-PD-L1 treated M1 macrophages in CD206 and Arg1 expression, the biological relevance of these changes was negligible when compared to TCM controls (**Supplementary Figure 2B**). When comparing PD-L1 KO and PD-L1 blockade TCM-polarized macrophages, no significant differences in fold change were observed in CD80 (**Figure 1H**), iNOS, CD206, or Arg1 expression (**Supplementary Figure 2C**). Thus, canonical inflammatory and tumor-conditioned markers, we did not observe broad differences in macrophage polarization caused by genetic deletion or antibody-mediated blockade of PD-L1 in vitro, especially compared with bona fide M1 or TCM marker levels. However, the decrease in CD80 surface levels upon either PD-L1 KO and blockade was consistent.

### PD-L1 blockade during and after macrophage polarization produces comparable effects

We tested the effects of PD-L1 blockade on maintenance of M1 and TCM polarization states by comparing M1 and TCM BMDMs treated with isotype or anti-PD-L1 antibody during a 96-hour window (**Figure 2A**, top row) with M1 and TCM BMDMs polarized for 48 hours prior to treatment with isotype or PD-L1 blockade for a subsequent 48 hours (**Figure 2A**, bottom row). Efficacy of PD-L1 blockade was confirmed by flow cytometry (**Supplementary Figure 3A**). The baseline isotype treated expression profile for each polarization condition was comparable between the control 96-hour polarization group and the experimental 48-hour treatment post 48-hour polarization group (**Supplementary Figure 3B-C**; “96hr Polarization” compared to “Post-Polarization”). Within M1 BMDMs in both conditions, CD80 surface levels were significantly reduced with anti-PD-L1 while iNOS did not change (**Figure 2B**). Analysis of TCM markers CD206 and Arg1 resulted in a significant decrease in CD206 expression in both conditions, but no significant change in Arg1 expression (**Figure 2C**). There was no significant difference in the log_2_ fold decrease in expression of CD80 and CD206 of PD-L1 blockade over isotype treated M1 BMDMs in both conditions (**Figure 2D**). While there was a statistically significant decrease and increase respectively in CD80 and iNOS levels in the 96-hour polarization group upon PD-L1 blockade, the expression levels were minimal compared to the positive controls **(Supplementary Figure 3D-E**).

**Figure 2:**
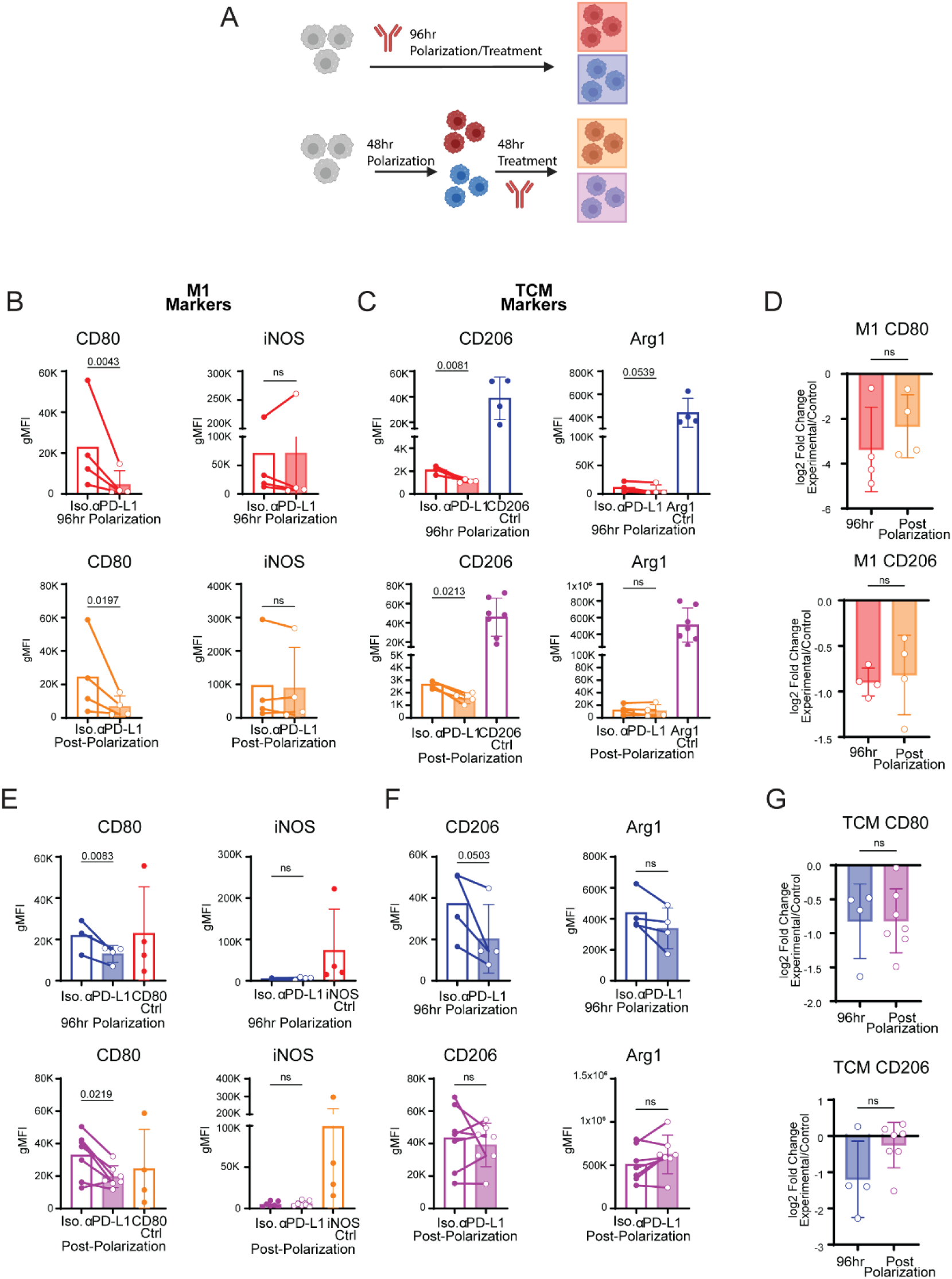
PD-L1 blockade during and after macrophage polarization produces comparable effects. (A) Experimental setup: Bone marrow cells isolated from C57BL/6J mice were differentiated into naïve M0 BMDMs using M-CSF, and M1 and TCM polarized by IFNy and LPS or IL-4/IL-10 and E0771 tumor media respectively. Macrophages were polarized for 96hrs. They were split into groups where they were either treated with anti-PD-L1 or Isotype control for 96hrs or allowed to polarize for 48hrs before being treated with antibodies for the second 48hrs. The 96hr polarization/treatment group was color coded Red=M1 and Blue=TCM. The Post-polarization treatment group was color coded Orange=M1 and Purple=TCM. (B) M1 marker expression of M1 polarized BMDMs 96hr polarization/treatment and post-polarization treatment (N=4 mice analyzed in 2 experiments)(CD80 geometric mean fold change 96hr = 0.14 (95% CI 0.06 – 0.31) post-polarization = 0.31 (95% CI 0.14 – 0.70)) (C) TCM marker expression of M1 polarized BMDMs 96hr polarization/treatment and post-polarization treatment (N=4 mice analyzed in 2 experiments)(CD206 geometric mean fold change 96hr = 0.54 (95% CI 0.40 – 0.74) post-polarization = 0.57 (95% CI 0.38 – 0.85)) (D) Log2 fold change of CD80 and CD206 expression comparing 96hr polarization/treatment with post-polarization treatment (N=4 mice analyzed in 2 experiments) (E) M1 marker expression of TCM polarized BMDMs 96hr polarization/treatment and post-polarization treatment (N=4-7 mice analyzed in 2-3 experiments)(CD80 geometric mean fold change 96hr = 0.59 (95% CI 0.45 – 0.77) post-polarization = 0.63 (95% CI 0.44 – 0.91)) (F) TCM marker expression of TCM polarized BMDMs 96hr polarization/treatment and post-polarization treatment (N=4-7 mice analyzed in 2-3 experiments) (CD206 geometric mean fold change 96hr = 0.48 (95% CI 0.23 – 1)) (G) Log2 fold change of CD80 and CD206 expression comparing 96hr polarization/treatment with post-polarization treatment (N=4-7 mice analyzed in 2-3 experiments) Data analysis by paired T test of log10 transformed data. Log2 fold change of Blockade over Isotype in 96hr and Post-Polarization groups performed per paired mouse and analyzed with Welch’s corrected T test. Constants 1499 and 2413 were used for M1 96hr and post-polarization CD206 samples. Constants 2196 and 2243.45 were used for TCM 96hr and post-polarization iNOS samples.

Within TCM-polarized BMDMs, the M1 marker CD80 was again significantly decreased in both PD-L1 blockade treatment during polarization and post-polarization while iNOS was unchanged (**Figure 2E**). The TCM marker CD206 trended towards a decrease in the 96-hour polarization treatment condition but not in the post-polarization condition, while Arg1 was unchanged in both (**Figure 2F**). When we compared the log_2_ fold changes in expression of CD80 and CD206 levels, we observed no difference in CD80 surface level fold changes between treatment conditions and a trend toward a larger decrease in CD206 surface levels during the 96-hour polarization condition (**Figure 2G**). Taken together, changes in polarization marker expression upon PD-L1 blockade post-polarization largely matched the changes observed upon treatment during polarization. Thus, we continued our analyses using PD-L1 blockade during polarization.

### Macrophage phagocytosis is unaffected by interference of PD-L1

We utilized pH-sensitive phRodo zymosan particles to determine general phagocytic capacity by imaging particle engulfment by BMDMs in vitro using the Incucyte live-cell imaging platform. Phagocytic rate was quantified as the area of phRodo fluorescence normalized to cell phase area over incubation time, with the area under the curve (AUC) calculated for WT versus PD-L1 KO (**Figures 3A-C**), and isotype versus PD-L1 blockade (**Figures 3D-F**) BMDMs in M0, M1, and TCM states. After 3 hours in culture, cells were harvested to quantify the total cell numbers and the frequencies of engulfed particles at the endpoint by flow cytometry. There were no significant changes to the phagocytic capacities of M0, M1, or TCM BMDMs in PD-L1 KO macrophages, as measured by Incucyte or by flow cytometric analysis (**Figures 3A-C**). There was also no change in phagocytic capacities of M0 and M1 BMDMs upon PD-L1 blockade (**Figures 3D-E**). While there was no change in particle engulfment by TCM BMDMs treated with PD-L1 blockade, there was a modest decrease in the percentage of phagocytic macrophages collected and analyzed by flow cytometry at the 3-hour timepoint (**Figure 3F**). Thus, functional macrophage activation, as reflected by zymosan engulfment, was not regulated by PD-L1 KO or blockade under the conditions tested.

**Figure 3:**
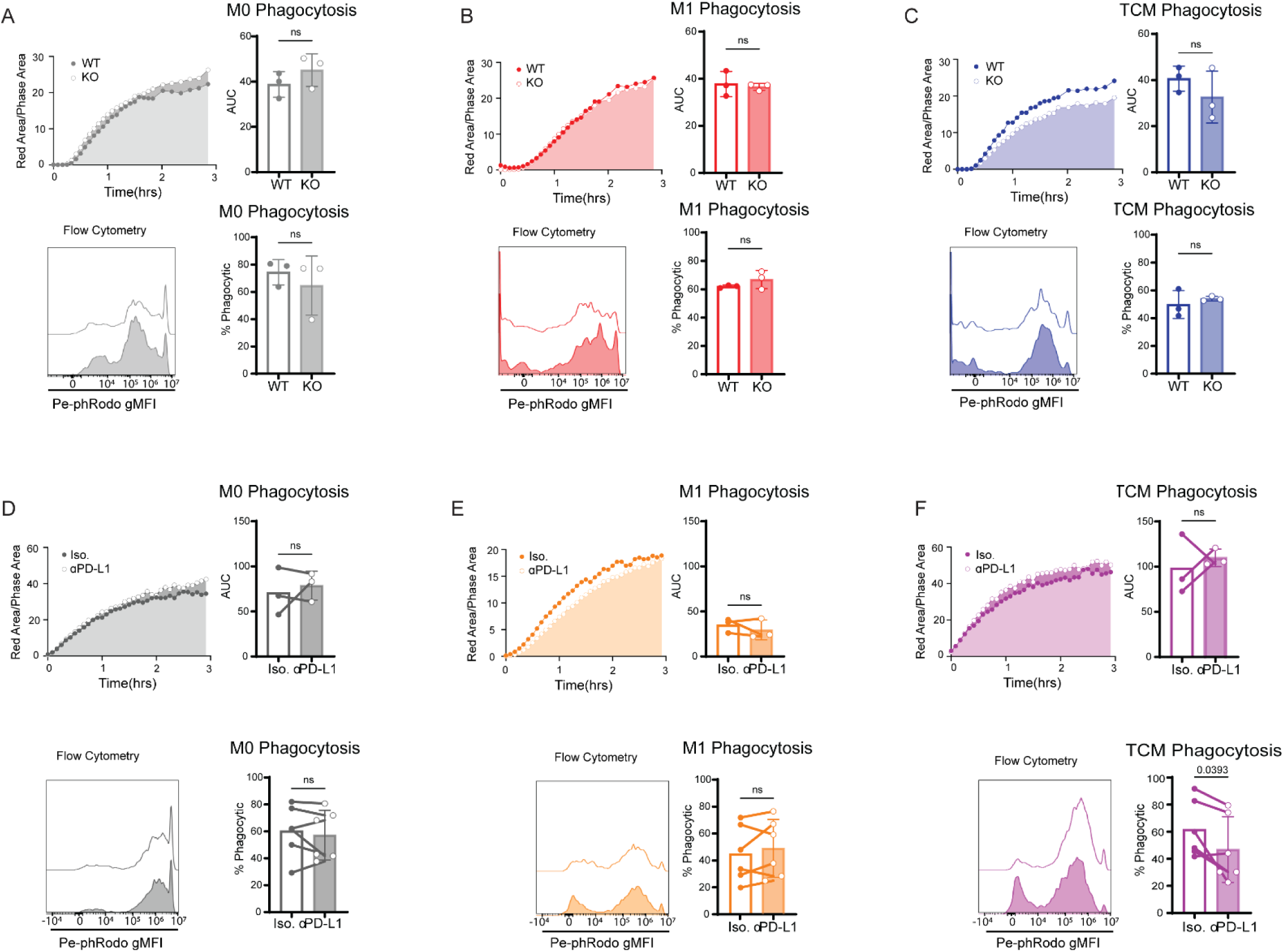
Macrophage phagocytosis is unaffected by interference of PD-L1. WT and KO BMDMs were incubated with pHrodo Red labeled zymosan particles and particle engulfment over time was measured, as defined by percentage of pHrodo-positive Red cell area normalized to total cell phase area. (A) Naïve BMDMs WT vs KO 3hr incubation summarized Red Area/Phase Area curves with corresponding analysis of the area under the curve (AUC) and flow cytometry analysis of red fluorescent BMDMs after the 3hr incubation. (N=3 Mice, each replicate a mean of triplicate wells) (B) M1 BMDMs WT vs KO AUC and flow analysis. (N=3 Mice, each replicate a mean of triplicate wells) (C) TCM WT vs KO AUC and flow analysis. (N=3 Mice, each replicate a mean of triplicate wells) (D) Naive Isotype vs anti-PD-L1 AUC and flow analysis. (N=3 Mice, each replicate a mean of 2-3 wells) (E) M1 Isotype vs anti-PD-L1 AUC and flow analysis. (N=3 Mice, each replicate a mean of 2-3 wells) (F) TCM Isotype vs anti-PD-L1 AUC and flow analysis. (N=3 Mice, each replicate a mean of 2-3 wells) Data analysis performed with Welch’s T test and paired T test for WT vs KO and Isotype vs anti-PD-L1 respectively.

### PD-L1 blockade and knockout differentially regulate antigen presentation and checkpoint molecule surface levels

Given the consistent reduction in CD80 surface levels across conditions, we expanded analysis to include antigen presentation molecules: major histocompatibility complexes class I (MHCI) and class II (MHCII), and immune checkpoint molecules: PD-1 and programmed cell death-ligand 2 (PD-L2). Similar to what was observed with polarization-associated molecules (CD80, iNOS, CD206, Arg1), WT M0 BMDMs and isotype control-treated BMDMs had low surface levels of MHCI, MHCII, PD-1, PD-L1, and PD-L2 (**Supplementary Figure 4**). Upon M1 polarization, WT BMDMs upregulated surface levels of MHCI, MHCII, and PD-L1, whereas WT TCM BMDMs upregulated MHCI, MHCII, PD-1, PD-L1, and PD-L2 (**Supplementary Figure 4A**). Isotype-treated WT BMDMs recapitulated these changes, with the exception of MHCII, in which TCMs expressed greater surface levels than M1 (**Supplementary Figure 4B**). Thus, M1 and TCM BMDMs have unique signatures with respect to antigen presentation machinery. WT and PD-L1 KO M0 BMDMs had similarly low surface levels of MHCI, MHCII, PD-1, and PD-L2 (**Supplementary Figure 4C**). PD-L1 blockade of M0 polarized BMDMs also did not impact surface levels of MHCI, MHCII, PD-1, or PD-L2 (**Supplementary Figure 4D**).

There were no changes to the surface levels of MHCI and MHCII between WT and PD-L1 KO M1 BMDMs, but PD-L1 blockade significantly decreased MHCII in M1 BMDMs (**Figure 4A**). Further analysis of M1 BMDMs indicated a statistically significant decrease in PD-1 surface levels in PD-L1 KO BMDMs as compared to WT, and a statistically significant increase in PD-L2 surface levels in PD-L1 blockade treated BMDMs as compared to isotype (**Figure 4B**). However, the biological significance of these changes in M1 BMDMs may be limited when compared to the 8-fold and 10-fold higher surface level expression of PD-1 and PD-L2 by TCM controls, respectively. PD-L1 KO TCMs showed significantly increased MHCII surface levels, approaching levels of MHCII-positive controls, whereas PD-L1 blockade significantly decreased MHCI but not MHCII surface levels compared to the isotype control (**Figure 4C**). Analysis of the checkpoint markers PD-1 and PD-L2, which are predominantly expressed in the TCM condition (**Supplementary Figure 4A**), demonstrated that PD-L1 KO BMDMs exhibited increased PD-1 surface levels while PD-L1 blockade decreased PD-1surface levels (**Figure 4D**). Additionally, in the TCM condition, PD-L2 surface levels were significantly decreased upon PD-L1 blockade as compared to isotype but was almost undetectable in PD-L1 KO (**Figure 4D**).

**Figure 4:**
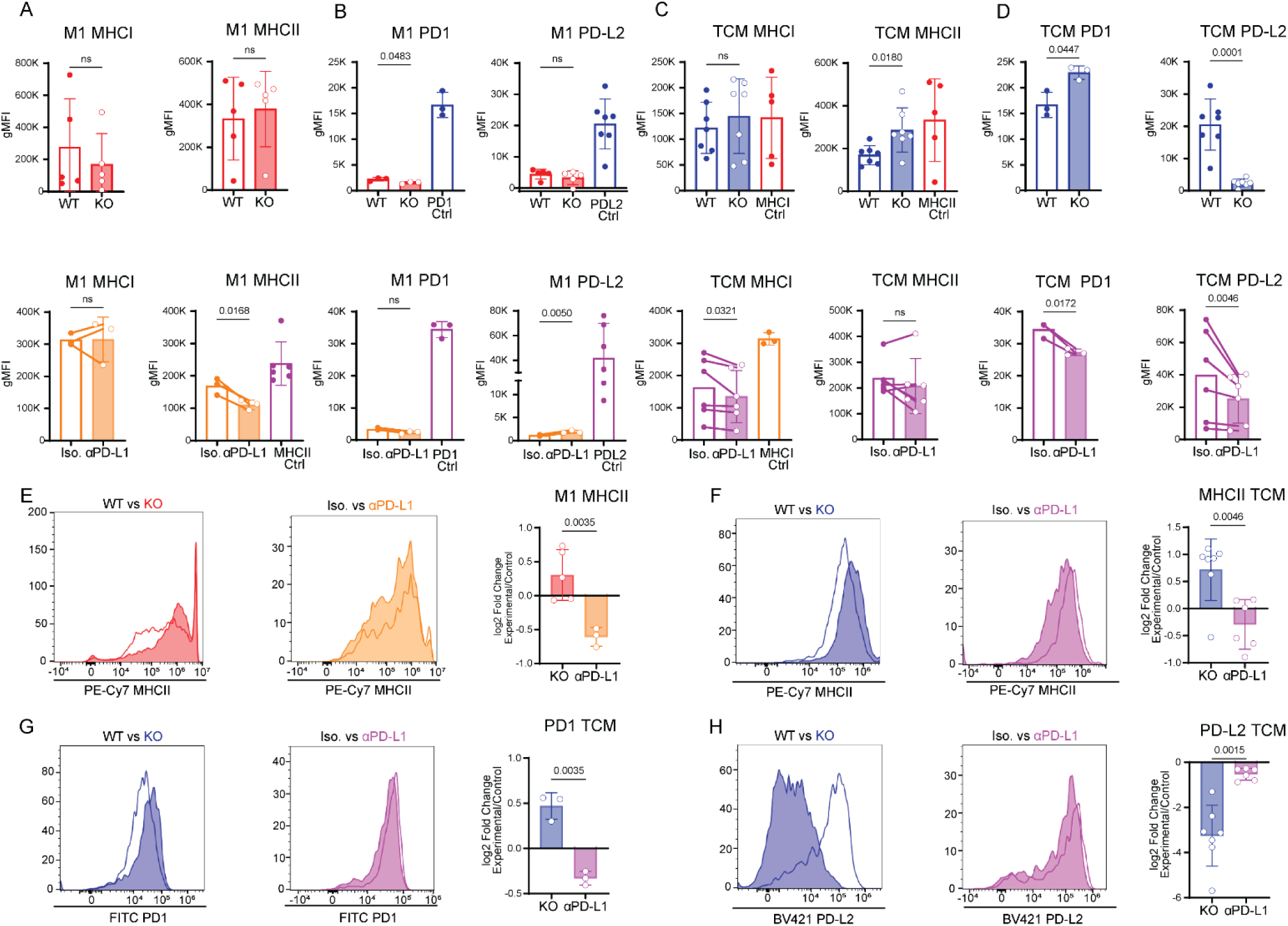
PD-L1 blockade and knockout differentially regulate antigen presentation and checkpoint molecules. (A) Antigen presentation molecules MHCI and MHCII expression in M1 polarized BMDMs comparing WT vs KO and Iso vs Blockade. (N=3-5 mice analyzed in 1-3 experiments) Geometric mean fold change MHCII = 0.66 (95% CI 0.52-0.83) (B) Checkpoint markers PD1 and PD-L2 in M1 polarized BMDMs comparing WT vs KO and Iso vs Blockade. (N=3-5 mice analyzed in 1-3 experiments) KO PD1 geometric mean fold change = 0.7 (95% CI 0.49-0.99) Blockade Geometric mean fold change PDL2 = 4.95 (95% CI 1.57 – 15.6)(C) Antigen presentation molecules MHCI and MHCII expression in TCM polarized BMDMs comparing WT vs KO and Iso vs Blockade. (N=7 mice analyzed in 5 experiments) KO Geometric mean fold changes: MHCII = 1.64 (95% CI 1.11 – 2.43) Blockade Geometric mean fold changes; MHCI = 0.82 (95% CI 0.69 – 0.98), (D) Checkpoint markers PD1 and PD-L2 in TCM polarized BMDMs comparing WT vs KO and Iso vs Blockade. (N=3-7 mice analyzed in 1-5 experiments). KO Geometric mean fold changes: PD1 = 1.38 (95% CI 1.02 – 1.89), PDL2 = 0.11 (95% CI 0.05 – 0.24). Blockade Geometric mean fold changes; PD1 = 0.8 (95% CI 0.7 – 0.91), PDL2 = 0.7 (95% CI 0.57 – 0.84). (E) Log2 fold change of KO/WT and blockade/iso of MHCII expression in M1 polarized BMDMs. (N=3-5 mice from 1-3 experiments) (F) Log2 fold change of KO/WT and blockade/iso of MHCII expression in TCM polarized BMDMs. (N=6-7 mice analyzed in 4-5 experiments) (G) Log2 fold change of KO/WT and blockade/iso of PD1 expression in TCM polarized BMDMs. (N=3 mice analyzed in 1 experiment) (H) Log2 fold change of KO/WT and blockade/iso of PD-L2 expression in TCM polarized BMDMs. (N=6-7 mice analyzed in 4-5 experiments) Analysis performed with Welch’s T test on log10 transformed data for WT vs KO groups. Paired T tests on log10 transformed data were used for Isotype vs Blockade groups. For both, constants to ensure positive values were used. Constants 1321 and 2021.67 for M1 PD-L2 WT/KO and Iso/aPD-L1 respectively. Welch’s T tests were performed on log2 fold change of experimental condition (KO, anti-PD-L1) to control condition (WT, Isotype) for PD-L1 deficiency and blockade BMDMs.

Comparing log_2_ fold changes in marker surface levels between PD-L1 KO and blockade revealed differential impact on MHCII surface levels, with PD-L1 KO exhibiting increased MHCII levels in both M1 (**Figure 4E**) and TCM conditions (**Figure 4F**), whereas MHCII levels were decreased in PD-L1 blockade treated M1 (**Figure 4E**) and TCM conditions (**Figure 4F**). long these lines, PD-1 surface levels were also differentially impacted, with an increase in PD-L1 KO TCM as compared to WT, and a decrease in PD-L1 blockade TCM as compared to isotype (**Figure 4G**). Comparison of log_2_ fold changes of PD-1 and PD-L2 surface levels in M1 polarized BMDMs revealed opposing effects of PD-L1 KO and blockade. PD-1 levels increased in PD-L1 KO macrophages but decreased upon PD-L1 blockade, whereas PD-L2 surface levels decreased upon PD-L1 KO and increased upon blockade. Analysis of TCM polarized BMDMs showed no significant differences in fold change in MHCI levels between PD-L1 KO and blockade groups (**Supplementary Figure 4E**). While PD-L2 levels were decreased in both PD-L1 KO and blockade groups, the magnitude of the decrease was significantly greater in PD-L1 KO TCM compared to WT than PD-L1 blockade TCM compared to isotype (**Figure 4H**). Together, these results indicate that PD-L1 KO and blockade have distinct effects on macrophage antigen presentation and checkpoint molecule surface levels, particularly in TCMs.

## Discussion

We investigated the cell-intrinsic role of PD-L1 in macrophages by comparing genetic deletion and antibody-mediated blockade with PD-L1-sufficient controls across multiple polarization states. Our findings suggest a macrophage intrinsic role of PD-L1 that influences the surface levels of immunomodulatory molecules on the cell surface. They also demonstrate that PD-L1 deletion and PD-L1 blockade differentially regulate macrophage immune phenotype, particularly surface levels of CD80, MHC molecules, and other T-cell interaction proteins. The distinct effects of the two tested methods of PD-L1 interference are consistent with PD-L1 having macrophage intrinsic functions beyond its known surface interactions.

Contrary to reports that PD-L1 blockade promotes inflammatory macrophage polarization, we observed minimal effects on classical polarization markers. Previous models indicated that PD-L1 blockade increased pro-inflammatory features MHCI, MHCII, and CD86 and decreased Arg1 by comparing fold-change of marker expression levels within macrophages treated with PD-L1 blockade and isotype controls(27,36). While these studies reported statistically significant changes in marker expression, the biological relevance of these results was less clear. In our analysis we compared the changes in marker expression with WT M1- or TCM-polarized macrophages as controls to provide context for the relative changes in polarization profiles. We also analyzed naïve, M1- and TCM-polarized macrophages, in which we demonstrated varied responses to PD-L1 interference. To ensure that the lack of changes in polarization was not due to preferential survival of less polarized macrophages upon polarization, we compared cell viability between WT, KO, isotype and antibody treated groups in each polarization condition and observed no decrease in viability (**Supplementary Figure 2**). Taken together, our findings suggest that PD-L1 perturbation is insufficient to broadly regulate macrophage polarization in isolated BMDMs. While perturbations to PD-L1 may induce statistically significant fold changes in polarization marker expression levels, they may not be so relevant when compared to that of bona fide M1 or TCM-polarized macrophages. However, this conclusion should be interpreted within the context of our in vitro system, as PD-L1 may exert more pronounced effects within tissue microenvironments where macrophages are exposed to complex cellular and inflammatory signals.

Although macrophage activation is frequently accompanied by changes in phagocytic function, neither genetic deletion nor antibody blockade of PD-L1 altered zymosan bead uptake in our system. Phagocytosis is a canonical function of macrophages that is frequently associated with macrophage activation. Prior studies have reported increased phagocytic function of macrophages following interference with the PD-1/PD-L1 axis in several tumor models, whereas in vitro monoculture systems have reported variable results(24,25,36,37). In our reductionist in vitro monoculture, neither genetic deletion nor antibody blockade of PD-L1 substantially altered zymosan particle uptake across polarization conditions. This finding is consistent with our observation that PD-L1 perturbation did not broadly alter macrophage polarization and suggests that the enhanced phagocytosis observed in vivo may depend on additional cellular and inflammatory cues in the tumor microenvironment rather than PD-L1 perturbation alone.

A consistent change to macrophage phenotype upon PD-L1 deletion or blockade was a reduction in CD80 surface levels, which we had also previously observed in our melanoma model (25). CD80 and PD-L1 have been shown to bind in cis on the plasma membrane of dendritic cells(38,39). Consistent with our findings, Zhao et al.(39) reported reduced CD80 surface levels following PD-L1 blockade and attributed this effect to CTLA4-mediated trans-endocytosis of CD80. However, because our experiments were performed in BMDM monocultures lacking CTLA4-expressing T cells, the consistent reduction in CD80 observed following both PD-L1 blockade and genetic deletion cannot readily be explained by this mechanism. Instead, our findings suggest that PD-L1 contributes to the maintenance of CD80 surface levels through a macrophage-intrinsic mechanism, potentially involving regulation of CD80 stability, trafficking, or surface retention.

The observed reduction in PD-L2 was particularly unexpected. Although direct interactions between PD-L1 and PD-L2 have not been described, both ligands bind PD-1 and share overlapping immunoregulatory functions. PD-L2 expression is typically lower than PD-L1, but exhibits approximately three-fold greater affinity for PD-1(40). In our model, PD-L1 was highly expressed in M1 macrophages and moderately expressed in TCMs, whereas PD-L2 surface expression was largely restricted to the TCM state (**Supplementary Figure 4**). Given these overlapping functions, we anticipated that loss of PD-L1 signaling might be accompanied by compensatory upregulation of PD-L2. Instead, PD-L2 surface levels decreased significantly following PD-L1 blockade and were nearly abolished in PD-L1-deficient macrophages (**Figure 4**). These findings suggest that PD-L1 and PD-L2 may be coordinately regulated rather than functionally compensatory, raising the possibility that PD-L1 contributes to the maintenance of PD-L2 surface levels. Further studies are needed to define the mechanisms underlying this relationship.

When comparing changes in MHCII surface levels, we observed opposing effects of PD-L1 genetic deletion and antibody-mediated blockade, with MHCII surface levels increasing following PD-L1 knockout but decreasing following PD-L1 blockade (**Figure 4**). This divergence is notable given the concurrent alterations in other molecules involved in T-cell communication, including CD80 and PD-L2. Together, our data indicated that genetic deletion and antibody-mediated blockade of PD-L1 do not produce equivalent effects on macrophage phenotype. These findings suggest that PD-L1 interference remodels the macrophage: T cell interaction landscape through coordinated changes in antigen presentation, co-stimulation, and checkpoint signaling pathways.

Although the specific effects of PD-L1 deletion and blockade differed across molecules, the most substantial changes were consistently observed in antigen presentation, co-stimulation, and checkpoint signaling. This is a surprising result when considering the prior literature suggested the largest changes would be seen in canonical polarization markers. These findings suggest that PD-L1 may have a greater influence on macrophage interactions with and instruction of adaptive immune cells than on canonical polarization profiles. One possible explanation for the disparity between our findings and previous studies is that PD-L1 dependent changes in antigen presentation, costimulatory, and checkpoint molecules alter macrophage interactions with nearby immune cells. These altered interactions could subsequently provide paracrine signals that influence changes in macrophage polarization and phagocytic capacity. In this model, macrophage-intrinsic PD-L1 functions would influence macrophage communication with the immune microenvironment, potentially exerting broader effects on macrophage activation only after reciprocal interactions with the tumor microenvironment. Such effects would not be expected in a reductionist monoculture system.

## Conclusions

The importance of macrophage PD-L1 in the tumor microenvironment continues to expand as the functions of both macrophages and PD-L1 are increasingly appreciated. Here, we demonstrate that genetic deletion and antibody-mediated blockade of PD-L1 produce distinct effects on macrophage phenotype. These findings support a cell-intrinsic role for PD-L1 in macrophages that extends beyond its canonical function as a ligand for PD-1 on T cells. Further investigation of the mechanisms by which PD-L1 regulates macrophage function may provide new opportunities to modulate the tumor microenvironment and improve the efficacy of immunotherapeutic strategies.

## Limitations

Our model was refined to one Tumor Conditioned Media E0771, while overlapping in some methodology with previous work done in the lab (Kwok et al., 2025), using the B16 melanoma model, some results differed. This suggests a highly tumor type dependent regulation of macrophage function. To further investigate PD-L1 functions in macrophages the ability to reintroduce PD-L1 expression in PD-L1 knockout backgrounds would add strength to these analyses. Because the PD-L1 KO model used is missing PD-L1 expression throughout maturation of monocytes to macrophages, there could be differences in macrophage maturation that would not be caused by blockade after differentiation. Reintroducing PD-L1 consructs could help parse out any differences due to PD-L1 loss before differentiation. Macrophage phagocytosis is regulated by diverse pathways which require multiple methods of analysis to clarify. Here we analyzed the effects of PD-L1 interference on a single phagocytic pathway. Our analysis does not include the effects of PD-L1 on phagocytosis of opsonized targets or apoptotic cells, but points to a general fitness in a core macrophage function.

## Future directions

The evidence we have gathered suggests that direct contact between PD-L1 and CD80 is important for maintaining CD80 expression levels on the cell surface. Previous studies have demonstrated the importance of PD-L1 and CD80 cis-interactions for dendritic cell migration and that the intracellular tail of PD-L1 is important for this interaction (39). However, it has not been clarified whether PD-L1 simply stabilizes CD80 on the cell surface or plays a role in its stabilization and trafficking to the cell surface. Further studies are needed to clarify this interaction. The loss of PD-L2 surface expression suggests a positive regulation of PD-L2 by PD-L1 expression. Due to the proximity of the PD-L2 gene to PD-L1, mechanisms of coregulation could contribute to the loss of PD-L2 upon PD-L1 knockout, but the decrease in PD-L2 upon PD-L1 blockade reduces the likelihood of that explanation. Investigating potential interactions of PD-L1 and upstream regulators of PD-L2 could clarify this dynamic.

## Materials and Methods

### Mice/Cell Culture

Wildtype C57Bl/6J mice were purchased from The Jackson Laboratories and PD-L1^-/-^ C57Bl/6 mice were generated and bred in-house(30). Mice were bred and maintained under specific-pathogen free conditions with controlled temperature and 12 hour light/dark day cycle in the Mayo Clinic animal facility. Female mice 6-10 weeks of age were used in experiments in accordance with protocols approved by the Mayo Clinic Institutional Animal Care and Use Committee (AUP #A00005857-21-R24).

E0771 murine triple negative breast cancer cells (ATCC #CRL-3461) were cultured under normoxic conditions at 37°C, 5% CO_2_ in Dulbecco’s Modified Eagle’s Medium (DMEM; Thermofisher), 10% fetal bovine serum (FBS; Thermofisher).

Naïve, M1, and TCM macrophages were treated with anti-PD-L1 (Clone 10F.9G2 BioXcell BE0101) or isotype matched (IgG2b LTF-2 BioXcell BE0090) at 20ug/mL during polarization (48hrs or 96hrs) and after polarization (48hrs) depending on experimental conditions.

### Generation of bone-marrow derived macrophages (BMDM)

BMDMs were generated using a modified version of the protocol described by (25,30). Bone marrow from the femur and tibiae of C57Bl/6J or PDL1^-/-^ C57Bl/6J mice were flushed out with serum free DMEM and cultured at densities between 2x10^6 and 4x10^6 per mL on 10cm, 6-well, or 12-well plates in DMEM supplemented with FBS (10%), Penicillin-Streptomycin (1%), and 20 ng/ml of M-CSF (Peprotech) for 6 days. Fresh media was replaced every 3 days. M0 macrophages were cultured in DMEM supplemented with M-CSF (20 ng/ml). Tumor-conditioned macrophages (TCMs) were generated by culturing macrophages in DMEM supplemented with M-CSF (20 ng/ml) and 20 ng/ml of IL-4 and IL-10 (Peprotech), and E0771 tumor conditioned media. M1 macrophages were generated by culturing macrophages in DMEM supplemented with M-CSF (20 ng/ml, Peprotech), 10 ng/ml of LPS (Sigma Aldrich), and 20 ng/ml of IFNγ (Peprotech). Macrophages were polarized for 48 hours then detached with Versene (Thermofisher), washed with PBS, then used for downstream applications such as flow cytometry, western blots, and imaging.

### Flow cytometry

In vitro cultures were washed with warm PBS, then detached from plates with Versene (Thermofisher) and washed with FWB (1x PBS + 2% fetal bovine serum). Cells were stained with fluorochrome-conjugated antibodies for 20min at 4°C, then washed with FWB. For intracellular staining, cells were fixed for 20 min at 4°C using the FOXP3 Fixation/Permeabilization kit (Thermofisher) as per manufacturer’s instructions then washed in permeabilization buffer. Cells were then stained with fluorochrome-conjugated intracellular antibodies for 20 min at 4°C, washed, and resuspended in FWB. Samples were read on a Cytek Aurora cytometer and data was analyzed using FlowJo software (v10, Treestar).

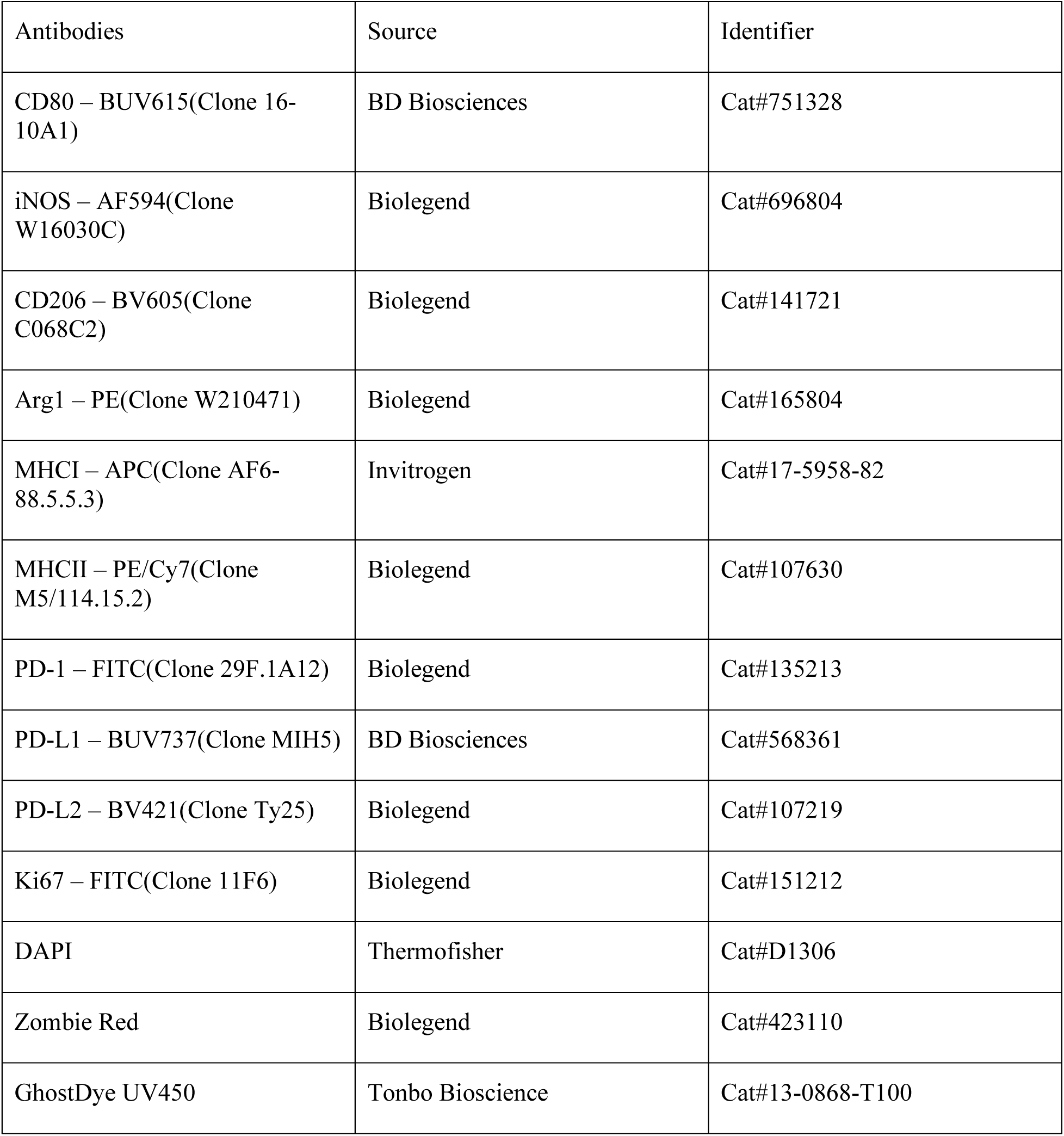

### Phagocytosis assays

pHrodo assays were performed with a modified protocol based on(25,42). BMDMs were polarized to M1 and TCM states for 24 hours, then washed and detached from wells using pre-warmed Trypsin EDTA for 5-10 minutes. Cells were counted and 5 X 10^4^ to 7.5 X 10^4^ were seeded per well into a 96-well flat bottom plate and polarization media returned to wells. After 24 more hours of incubation, 2.5ug of red Zymosan pHrodo bioparticles (Sartorius) were added to each well and placed in an Incucyte incubator (Sartorius), scanned every 5-10 minutes for 3 hours at 10x magnification. After the 3 hour analysis with the incucyte, cells were washed and detached for flow cytometry analysis.

### Cell Cycle/Viability

2.5 x 10^6^ Bone marrow cells were plated and differentiated in 6-well plates. Once differentiated, they were polarized to M1 and TCM states and WT BMDMs treated with anti-PD-L1 antibody or Isotype controls. Cells were detached using trypsin for 5-10 minutes and stained for viability. They were then fixed using the FOXP3 Fixation/Permeabilization kit and stained for Ki67 for 20min at 4°C. They were then washed and stained with DAPI for 3-5 minutes at room temperature. They were then analyzed using a BD Symphony cytometer.

### Western Blot

Whole cell lysates were prepared in RIPA buffer containing Halt protease inhibitors. Equal amounts of protein (30ug) were separated by SDS-PAGE, transferred to low fluorescence PVDF membranes using a turbotransfer system, and probed with antibodies against PD-L1 (MIH5 eBioscience 14-5982-82) and GAPDH(14C10 Cell Signaling 2118). Blots were visualized using IRDye conjugated secondary antibodies using a LI-COR Odyssey Clx imaging system.

### Statistical Analysis

Data in this study are presented as means ± SEM for n = 3-7 representing the number of mice in each assay. Polarization controls were not included in the analysis. Statistical analysis was performed using GraphPad Prism software and can be found in the corresponding figures and figure legends. Depending on the data, Welch’s T test, Lognormal Welch’s T test, Paired and Ratio paired T tests, one-way ANOVA, mixed effects and lognormal one-way ANOVA were performed.

## Author Contributions

T.W. and J.L. designed the experiments. T.W. performed the experiments and analyzed the data. M.R.-J. and H.D. provided input for research design and interpretation. T.W. and J.L. wrote the manuscript. T.W., M.R.-J., H.D., and J.L. reviewed and edited the manuscript.

## Acknowledgments

We thank the Mayo Clinic Arizona Flow Cytometry Core for experimental and technical support. We thank Dr. Ildefonso Silva-Junior, Caio Loureiro Salgado for technical support in flow cytometry assays. We thank Dr. Tina Kwok for technical support in experimental protocols and early conceptual work. And we thank the laboratory of Dr. William Faubion (Mayo Clinic, Phoenix, AZ, USA) for support and equipment for western blot assays.

## Supplementary Data

**Supplementary Figure 1.**
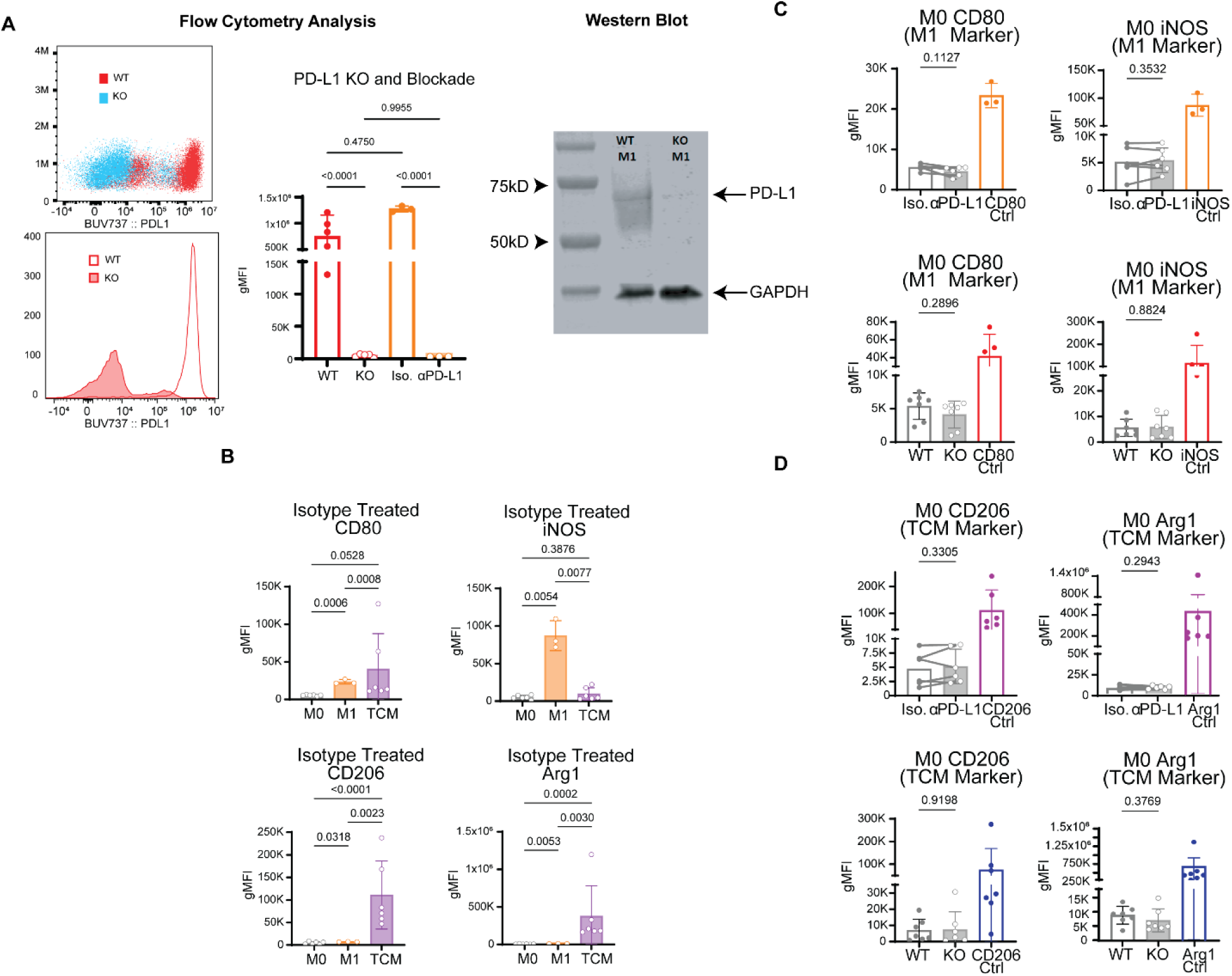
(A) PD-L1 knockout and blockade confirmation through flow cytometry (left figure) and western blot confirmation of PD-L1 knockout. All M1 polarized. Demonstrating compete knockout and complete blockade of surface PD-L1. Statistical analysis was performed using a Lognormal ordinary one-way ANOVA with Tukey’s multiple comparisons Test. (B) Baseline expression of polarization panel in Isotype treated BMDMs. CD80 TCM expression appears higher than M1 BMDMs due to experimental day variability resulting in outliers. Experimental day matched samples result in M1 BMDMs with the highest expression. Statistical analysis was performed using Mixed-effects analysis with Tukey’s multiple comparisons. (C) M1 marker expression of naïve BMDMs WT/KO and Iso/aPD-L1. (D) TCM marker expression of naïve BMDMs WT/KO and Iso/aPD-L1. The remaining statistical analysis was performed with Welch’s T test analysis of Log10 transformed data for WT/KO analysis and Ratio paired T test for Iso/aPD-L1 analysis.

**Supplementary Figure 2.**
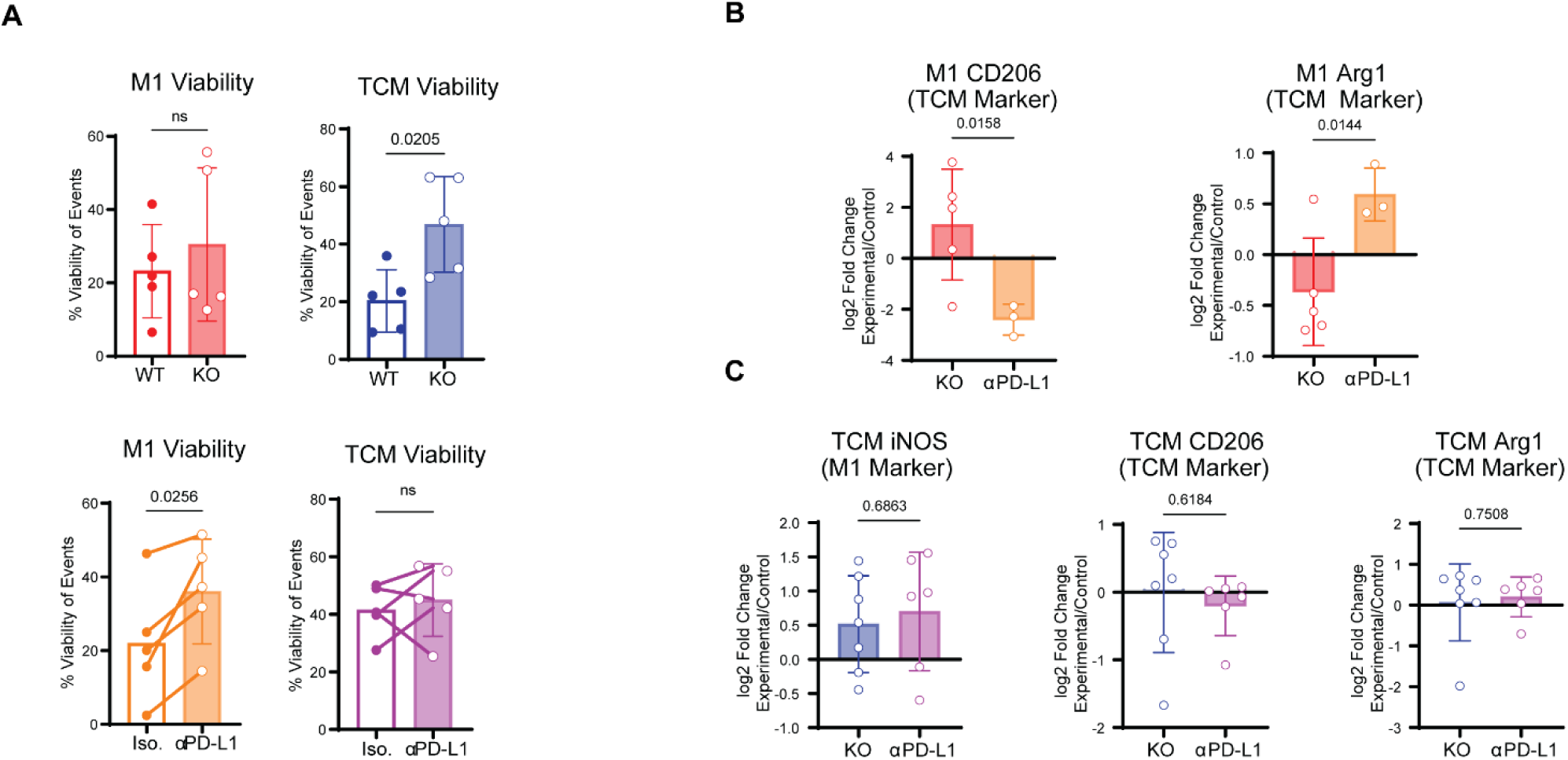
(A) Cell viability analyzed using a viability dye as the fraction of viable cells of the total cell population lifted from the well. (N=5 mice from 1 experiment) Statistical analysis performed using Welch’s T test for WT/KO and Paired T test for Iso/aPD-L1. (B) Log2 fold change of KO/WT and aPD-L1/iso of CD206 and Arg1 expression in M1 polarized BMDMs. (N=3-5 mice from 1-3 experiments) Statistical analysis performed using Welch’s T test. (C) Log2 fold change of KO/WT and aPD-L1/iso of iNOS, CD206, and Arg1 expression in TCM polarized BMDMs. (N=6-7 mice from 3-5 experiments) Statistical analysis performed using Welch’s T test.

**Supplementary Figure 3.**
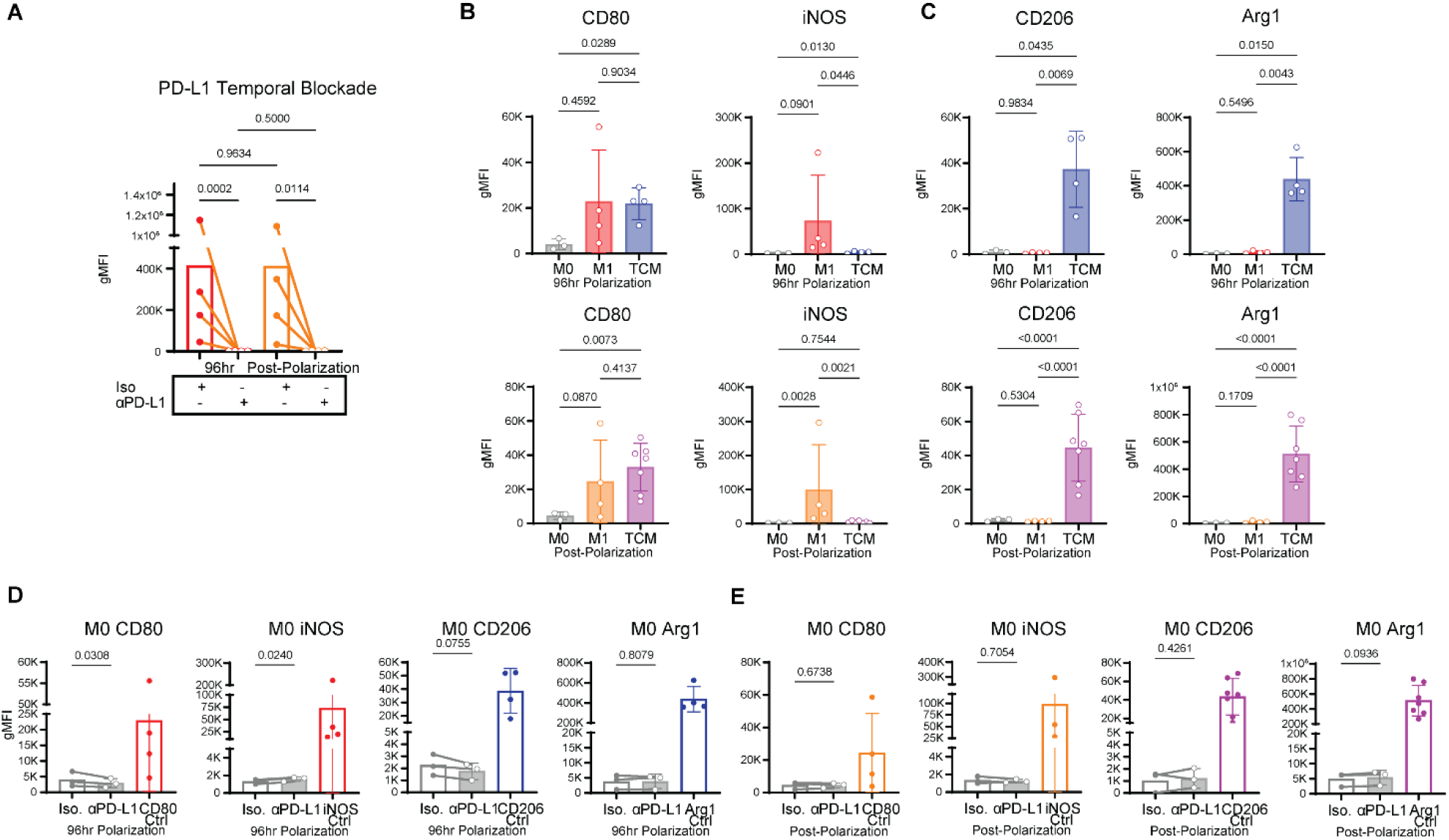
(A) Flow cytometry analysis performed to confirm blockade of PD-L1 in the 96hr and post-polarization M1 polarized treatment groups. (N=4 mice from 2 experiments) Statistical analysis was performed using a Lognormal ordinary one-way ANOVA with Tukey’s multiple comparisons Test. (B) Baseline expression of M1 polarization markers in 96hr and post-polarization groups. (N=3-7 mice from 1-4 experiments.) Statistical analysis was performed using Mixed-effects analysis with Tukey’s multiple comparisons. (C) Baseline expression of TCM polarization markers in 96hr and post-polarization groups. (N=3-7 mice from 1-4 experiments.) Statistical analysis was performed using Mixed-effects analysis with Tukey’s multiple comparisons. (D) Polarization marker expression of naïve BMDMs in the 96hr treatment group. (E) Polarization marker expression of naïve BMDMs in the post-polarization treatment group. The remaining statistical analysis was performed with Ratio paired T test.

**Supplementary Figure 4.**
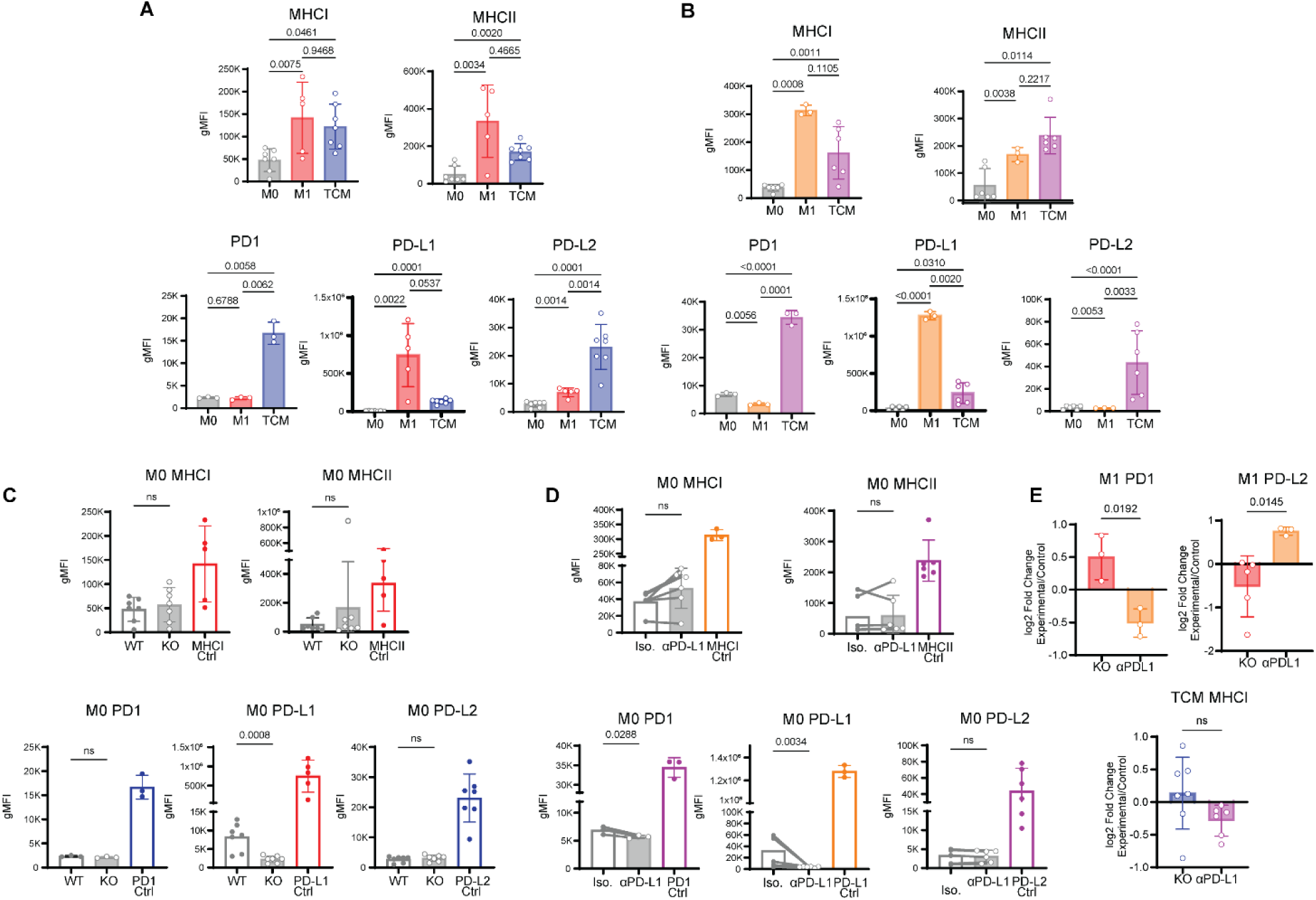
(A) Baseline expression of antigen presentation and checkpoint markers of WT BMDMs. (N=3-7 from 1-4 experiments.) Statistical analysis was performed using Mixed-effects analysis with Tukey’s multiple comparisons. (B) Baseline expression of antigen presentation and checkpoint markers of Isotype treated BMDMs. (N=3-6 from 1-4 experiments.) Statistical analysis was performed using Mixed-effects analysis with Tukey’s multiple comparisons. (C) Antigen presentation and checkpoint marker expression of WT/KO naïve BMDMs. (N=3-7 from 1-4 experiments). (D) Antigen presentation and checkpoint marker expression of Iso/aPD-L1 naïve BMDMs. (N=3-6 from 1-4 experiments). C and D statistical analysis performed with Welch’s T test analysis of Log10 transformed data for WT/KO analysis and Ratio paired T test for Iso/aPD-L1 analysis. (E) Log2 fold change of KO/WT and aPD-L1/iso of PD1 and PD-L2 expression in M1 polarized BMDMs. (N=3-5 mice from 1-3 experiments) and MHCI expression in TCM polarized BMDMs. (N=5-7 mice from 3-4 experiments) Statistical analysis performed using Welch’s T test.

